# High-resolution genome-wide functional dissection of transcriptional regulatory regions in human

**DOI:** 10.1101/193136

**Authors:** Xinchen Wang, Liang He, Sarah Goggin, Alham Saadat, Li Wang, Melina Claussnitzer, Manolis Kellis

## Abstract

Genome-wide epigenomic maps revealed millions of regions showing signatures of enhancers, promoters, and other gene-regulatory elements^1^. However, high-throughput experimental validation of their function and high-resolution dissection of their driver nucleotides remain limited in their scale and length of regions tested. Here, we present a new method, HiDRA (High-Definition Reporter Assay), that overcomes these limitations by combining components of Sharpr-MPRA^2^ and STARR-Seq^3^ with genome-wide selection of accessible regions from ATAC-Seq^4^. We used HiDRA to test ~7 million DNA fragments preferentially selected from accessible chromatin in the GM12878 lymphoblastoid cell line. By design, accessibility-selected fragments were highly overlapping (up to 370 per region), enabling us to pinpoint driver regulatory nucleotides by exploiting subtle differences in reporter activity between partially-overlapping fragments, using a new machine learning model SHARPR2. Our resulting maps include ~65,000 regions showing significant enhancer function and enriched for endogenous active histone marks (including H3K9ac, H3K27ac), regulatory sequence motifs, and regions bound by immune regulators. Within them, we discover ~13,000 high-resolution driver elements enriched for regulatory motifs and evolutionarily-conserved nucleotides, and help predict causal genetic variants underlying disease from genome-wide association studies. Overall, HiDRA provides a general, scalable, high-throughput, and high-resolution approach for experimental dissection of regulatory regions and driver nucleotides in the context of human biology and disease.

## Introduction

Precise spatiotemporal control of gene expression is achieved by the interplay between non-coding regulatory elements, including distal enhancers and proximal promoters, and the transcriptional regulators they help recruit or repel, thus modulating the expression of nearby genes^5,6^. Unlike protein-coding genes, which can be readily identified by their sequence properties and evolutionary signatures, gene-regulatory elements lack highly-predictive sequence patterns and show only modest evolutionary conservation at the nucleotide level^5,7^. Thus, systematic recognition of gene-regulatory elements has relied on mapping of their epigenomic signatures, including DNA accessibility, histone modifications, and DNA methylation^1,8,9^. For example, both enhancers and promoters have high DNA accessibility and low H3K27me3, but distal enhancers show relatively higher H3K27ac and H3K4me1 while promoters show relatively higher H3K9ac and H3K4me3^10,11^. However, many regions showing such epigenomic marks do not experimentally drive reporter gene expression, and some regions driving gene expression lack endogenous signatures^12-14^.Moreover, epigenomic signatures are often low-resolution, with important driver regulatory nucleotides comprising only a small subset of the larger regions showing epigenomic signatures^2^.

Experimental dissection of enhancer and promoter regions has been traditionally expensive, laborious, low-throughput, and low-resolution, lacking the resolution to pinpoint individual regulatory driver nucleotides without recourse to extensive mutagenesis. Several high-throughput reporter assays for enhancer function have recently been developed, enabling the testing of tens of thousands of distinct DNA sequences simultaneously, including MPRA and CRE-Seq^15-17^. These use microarray-based oligonucleotide synthesis technology to generated the tested elements and their barcodes, clone them into a common episomal reporter vector, and use high-throughput sequencing to quantify expression. Technical limitations of oligonucleotide synthesis currently restrict the maximum tested DNA fragment length to ~230 nucleotides, and the maximum number of tested constructs to ~240,000 sequences per array. Although still limited in the number of target regions, Sharpr-MPRA enabled higher-resolution inferences by densely tiling target regions with multiple overlapping constructs, and exploiting subtle differences between the measured activity of neighboring constructs to achieve offset resolution (~5bp) instead of construct resolution (~230bp)^2^. STARR-Seq integrated random genomic fragments downstream of the transcription start site of episomal reporter genes, thus foregoing the oligo synthesis step and the need for barcodes as the tested elements were transcribed and serve as their own activity reporters^3^. However, STARR-seq fragments are selected by random genomic fragmentation. As random genomic fragmentation does not densely cover regulatory elements, STARR-seq has limited efficiency and resolution at regulatory regions.

Here, we present HiDRA, a novel high-resolution global screen for transcriptional regulatory activity in accessible regions, building on several key ideas from previous technologies to overcome their limitations and combine their advantages, enabling high-efficiency, high-throughput, and high-resolution inference of regulatory activity. We extract accessible DNA regions from ATAC-Seq^4^, size-select for constructs 150-500nt long, and insert them downstream of episomal reporters genes to test their activity and exploit their overlapping nature for high-resolution inferences. Our approach overcomes the construct-length and region-count limitations of synthesis-based technologies, and our ATAC-seq selection of open chromatin regions concentrates the signal on likely regulatory regions and enables high-resolution inferences. Altogether, we test enhancer constructs of comparable length to low-throughput studies, achieve the high resolution dissection of systematic perturbation studies, and test millions of unique fragments in a single experiment.

We applied HiDRA to infer genome-wide regulatory activity across ~7 million DNA fragments preferentially selected from accessible chromatin in the GM12878 lymphoblastoid cell line, resulting in ~95,000 active fragments clustering in ~65,000 regions showing significant regulatory function. These are enriched for endogenous active histone marks (including H3K9ac, H3K27ac), regulatory sequence motifs, and regions bound by immune regulators. Our ATAC-based selection approach resulted in highly-overlapping fragments, with up to 370 fragments per region, enabling us to pinpoint driver regulatory nucleotides. We discover ~13,000 high-resolution driver elements, which are enriched for regulatory motifs and evolutionarily conserved nucleotides, and help predict causal genetic variants underlying disease from genome-wide association studies. Overall, HiDRA provides a general, scalable, high-throughput, and high-resolution approach for experimental dissection of regulatory regions and driver nucleotides in the context of human biology and disease.

## Results

### HiDRA experimental method overview and plasmid library construction

HiDRA leverages the selective fragmentation of genomic DNA at regions of open chromatin to generate fragment libraries that densely cover putative transcriptional regulatory elements. Fragments are enriched from open chromatin and regulatory regions using ATAC-seq (Assay for Transposase-Accessible Chromatin with high throughput sequencing) and subsequently cloned into the 3’ untranslated region (UTR) of a reporter gene on the self-transcribing enhancer reporter vector used in STARR-seq^3,4^. Fragments with transcriptional regulatory activity promote self-transcription such that active segments of DNA can be identified and quantified by high-throughput RNA sequencing to produce a quantitative readout of enhancer activity (**Fig. 1a**). Library construction can be completed in 2-3 days and requires as few as 10^4^-10^5^ cells as input starting material.

**Figure 1:**
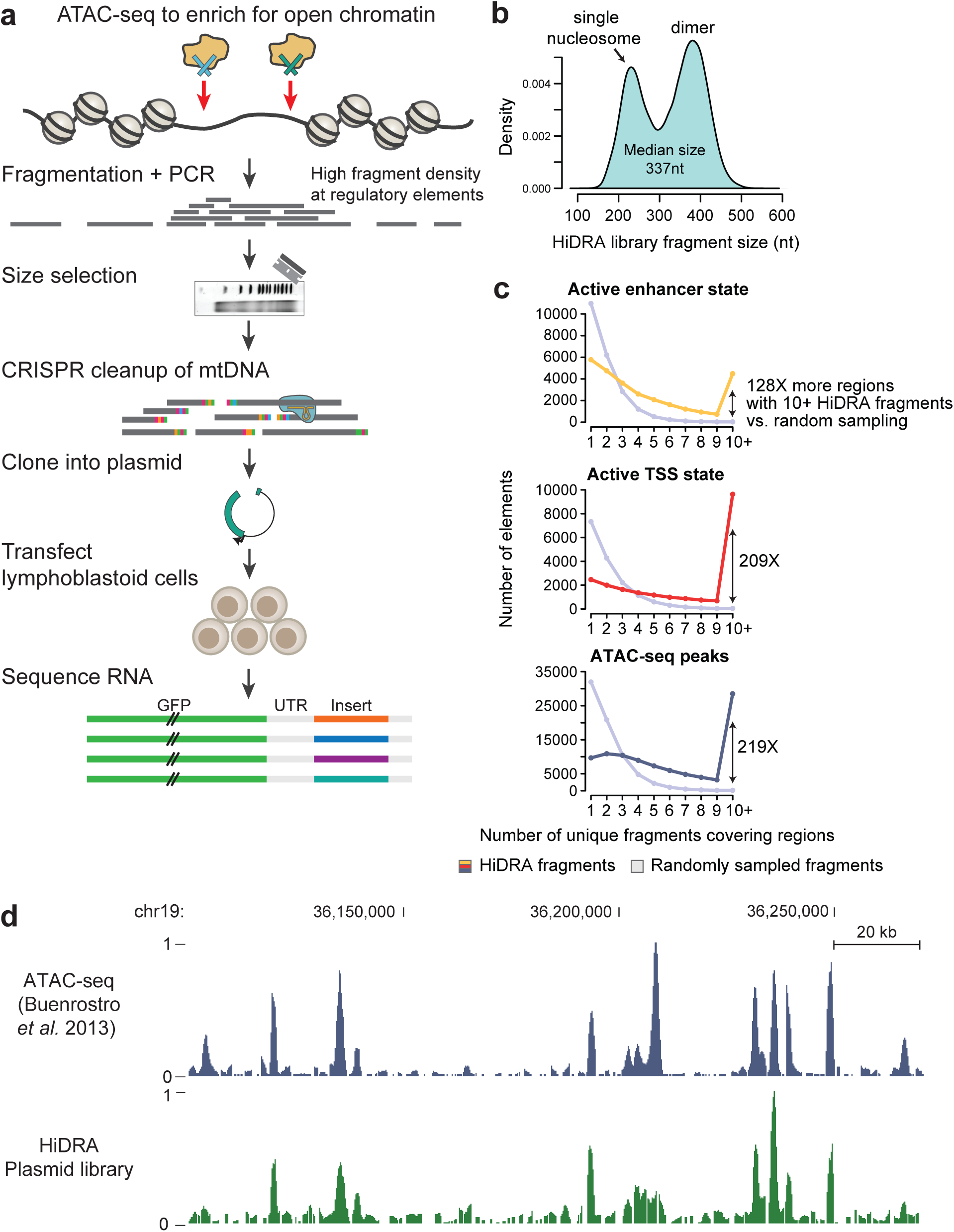
Overview of HiDRA library preparation. (a) The Tn5 transposase preferentially fragments genomic DNA at regions of open chromatin. Fragments are then size-selected on an agarose gel and mtDNA contamination is removed by selective CRISPR-Cas9 degradation. The fragment library is amplified by PCR and cloned into a enhancer reporter vector. (b) Size distribution of HiDRA library fragments. Bimodal shape is due to Tn5 preference to cut adjacent to nucleosomes (c) Number of predicted enhancer, active TSS and ATAC-seq peaks covered by multiple unique HiDRA fragments. (d) HiDRA plasmid library recapitulates genomic coverage of a conventional ATAC-seq experiment.

We constructed a HiDRA library with 9.7 million total unique mapping fragments, of which 4 million had a frequency greater than 0.1 reads per million (RPM; non-mitochondrial reads). More than 99% of fragments had lengths between 169 nt and 477nt (median: 337nt), with the fragment length distribution showing two peaks spaced by ~147 nt, corresponding to the length of DNA wrapped around each nucleosome (**Fig. 1b**). In contrast to unbiased fragmentation of the genome, our library has much higher efficiency for selectively targeting accessible DNA regions that are more likely to play gene-regulatory roles. Our HiDRA library covers 4486 predicted enhancers and 9631 predicted promoters (“Active Transcription Start Site (TSS)” state) with more than 10 unique fragments (**Fig. 1c**, colored lines), a ~130-fold and ~210-fold enrichment compared to 35 enhancer and 46 promoter regions expected to be covered by chance at the same coverage, indicating that HiDRA library construction successfully targets predicted regulatory regions. Even among enhancer and promoter regions, those with higher expected activity are preferentially selected by HiDRA, as they show higher accessibility and are thus more likely to be cloned in our library and tested by our episomal reporters (Supplemental **Fig. S1**).

Our cloning strategy is specifically designed to densely sample regulatory regions, in order to enable high-resolution inference of regulatory activity from highly-overlapping fragments. Indeed, we found up to 370 unique fragments per region in our HiDRA libraries, with ~32,000 genomic intervals containing at least 10 overlapping fragments and ~2750 containing at least 50 fragments, compared to 180 and 0 that would be expected by randomly-selected fragments, respectively. In addition to clustering of tested fragments within the same region, high-resolution inference relies on partially-overlapping rather than fully-overlapping fragments, which requires a random fragmentation pattern. Indeed, the Tn5 transposase we used here inserts randomly into the genome, and indeed the resulting DNA fragments provide a dense sampling of start and end positions that mirrors the peaks of ATAC-seq experiments (**Fig. 1d**), indicating that accessible regions most likely to show regulatory activity will have both higher representation in our libraries, and also more starting and ending positions that can help identify driver nucleotides.

### Identification of DNA fragments with transcriptional regulatory activity

To evaluate the ability of each cloned DNA fragment to promote gene expression, we transfected our HiDRA library into GM12878 lymphoblastoid cells, collected RNA 24 hours post-transfection, and measured the abundance of transcribed fragments by high-throughput RNA sequencing. We carried out 5 replicate transfection experiments from the same plasmid library, each into ~120 million cells, and we observed a high degree of correlation in the RNA counts between replicates (0.95 Pearson correlation on average for fragments ≥1 RPM; 0.76 for ≥1 RPM; Supplemental **Fig. S2**). To quantify the regulatory activity of tested elements, we compared the number of RNA reads obtained for a fragment (corresponding to the expression level of the reporter gene, as the constructs are self-transcribing), relative to representation of that fragment in the non-transfected input plasmid library (thus normalizing the differential abundance of each fragment in our library). We observed a substantial number of fragments that are more prevalent in RNA than DNA, indicating capability of many HiDRA fragments to drive reporter gene expression (**Fig. 2a**).

**Figure 2:**
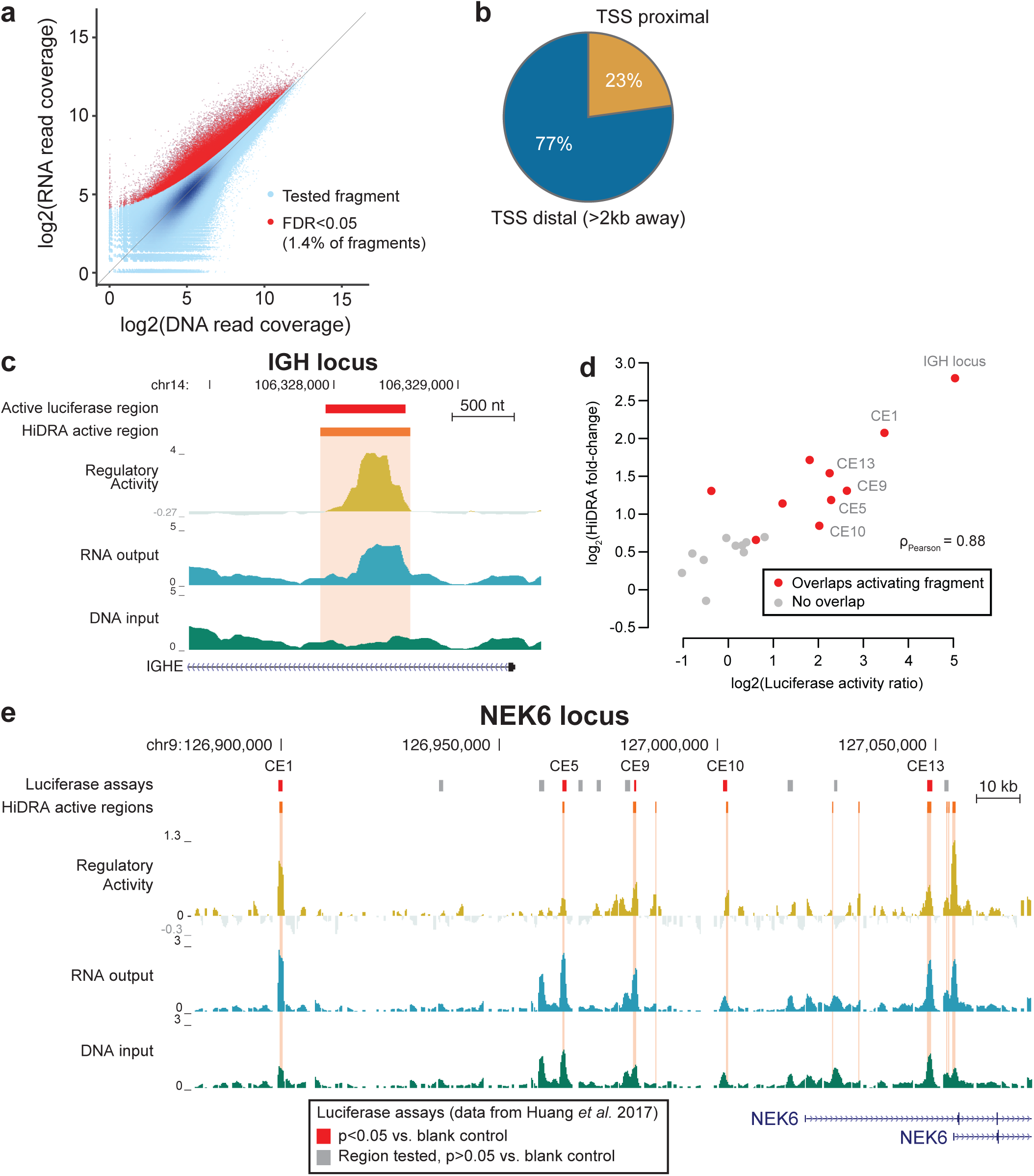
HiDRA identifies transcriptional regulatory elements. (a) Scatterplot of abundances for HiDRA fragments in input plasmid DNA) and output (RNA) samples. Abundances calculated after merging all five replicates. Active HiDRA fragments alled by DESeq2 highlighted with red dots, blue color intensity corresponds to greater density of points. (b) The majority of HiDRA active regions are distal to annotated TSSs (>2kb). (c) HiDRA identifies enhancer activity within an intron in the immu-oglobulin heavy chain locus. Red bar, DNA segment active in luciferase assay performed by Huang *et al.* (2017). Orange bar nd highlight, region identified by HiDRA as having transcriptional regulatory activity. (d) Quantitative comparison of luciferase ssay activity levels to HiDRA for 21 predicted enhancer elements. HiDRA signal corresponds to maximum activity within the egion tested by luciferase, and luciferase value corresponds to median normalized activity over biological replicates. Pearson orrelation calculated after log2 transformation. (e) Comparison of HiDRA-called active regions with luciferase assay results for 3 enhancers at the NEK6 locus. Luciferase experiments are colored in red or grey depending on whether DNA fragmentsrive luciferase activity in GM12878 cells as determined by Huang *et al.* (2017).

Given the intentionally high initial complexity of our HiDRA library, many fragments will be sequenced with a relatively low depth of coverage. We therefore grouped fragments with a 75% reciprocal overlap to boost the read coverage of genomic regions and increase statistical power. This yielded 7.1 million unique “fragment groups” generated from merging 9.7 million HiDRA fragments. In total, we identified 95,481 fragment groups that promote reporter gene expression at an FDR cut-off of 0.05, which we will refer to as ‘active HiDRA fragments’ (**Fig. 2a**, red dots). These 95,481 active HiDRA fragments are located within 66,254 unique genomic intervals that we subsequently refer to as “active HiDRA regions”.

We found that active HiDRA fragments showed a wide range of input DNA levels in our plasmid library, indicating that regulatory function and DNA accessibility rely on complementary sequence signals, and that DNA accessibility alone is not sufficient to predict episomal regulatory function. We also found that active HiDRA regions are predominantly distal to annotated transcription start sites (TSSs) (**Fig. 2b**), validating the utility of HiDRA for pinpointing distal regulatory regions that are particularly challenging to identify.

As proof-of-concept that HiDRA is capable of identifying true enhancer elements, we examined the well-studied immunoglobulin heavy chain enhancer within the intron of the immunoglobulin heavy constant epsilon (IGHE) gene^18^. We observed that the peak of HiDRA activity is centered precisely within the region previously identified as driving enhancer activity in low-throughput luciferase assays (**Fig. 2c**).

To assess the quantitative accuracy of HiDRA relative to luciferase assays, we compared active HiDRA regions and luciferase results across 21 putative enhancers predicted and tested independently by Huang *et al.*^19^. We found a 0.88 Pearson correlation between measured luciferase activity and HiDRA activity, confirming the accuracy and quantitative nature of our high-throughput approach (**Fig. 2d**). A visualization of 14 luciferase-tested enhancers in the serine/threonine kinase *NEK6* locus shows a strong correspondence between luciferase assay results and HiDRA active regions (**Fig. 2e**).

### HiDRA regulatory elements are enriched in promoter and enhancer elements

We next surveyed the 95,481 active HiDRA fragments identified in GM12878 to assess shared common genomic or epigenomic characteristics. In comparison to the set of all HiDRA fragments tested, active fragments were 12 times more likely to overlap an active promoter “TssA” chromatin state (marked by H3K4me3 and H3K27ac, **Fig. 3a** *inset*) and 5 times more likely to overlap an “Active Enhancer” chromatin state (marked by H3K4me1 and H3K27ac, **Fig. 3a**). By contrast, “Weak Enhancer” chromatin states (marked by H3K4me1 but lack of H3K27ac) showed substantially weaker enrichment (2.2-fold) within active HiDRA fragments than active enhancers, consistent with previous literature indicating that presence of H3K27ac correlates with higher greater expression of nearby genes (**Fig. 3b**). Overall, 35% of all predicted active promoters (8355 regions) and 16% of all predicted active enhancers (5276 regions) overlapped at least one active HiDRA fragment.

**Figure 3:**
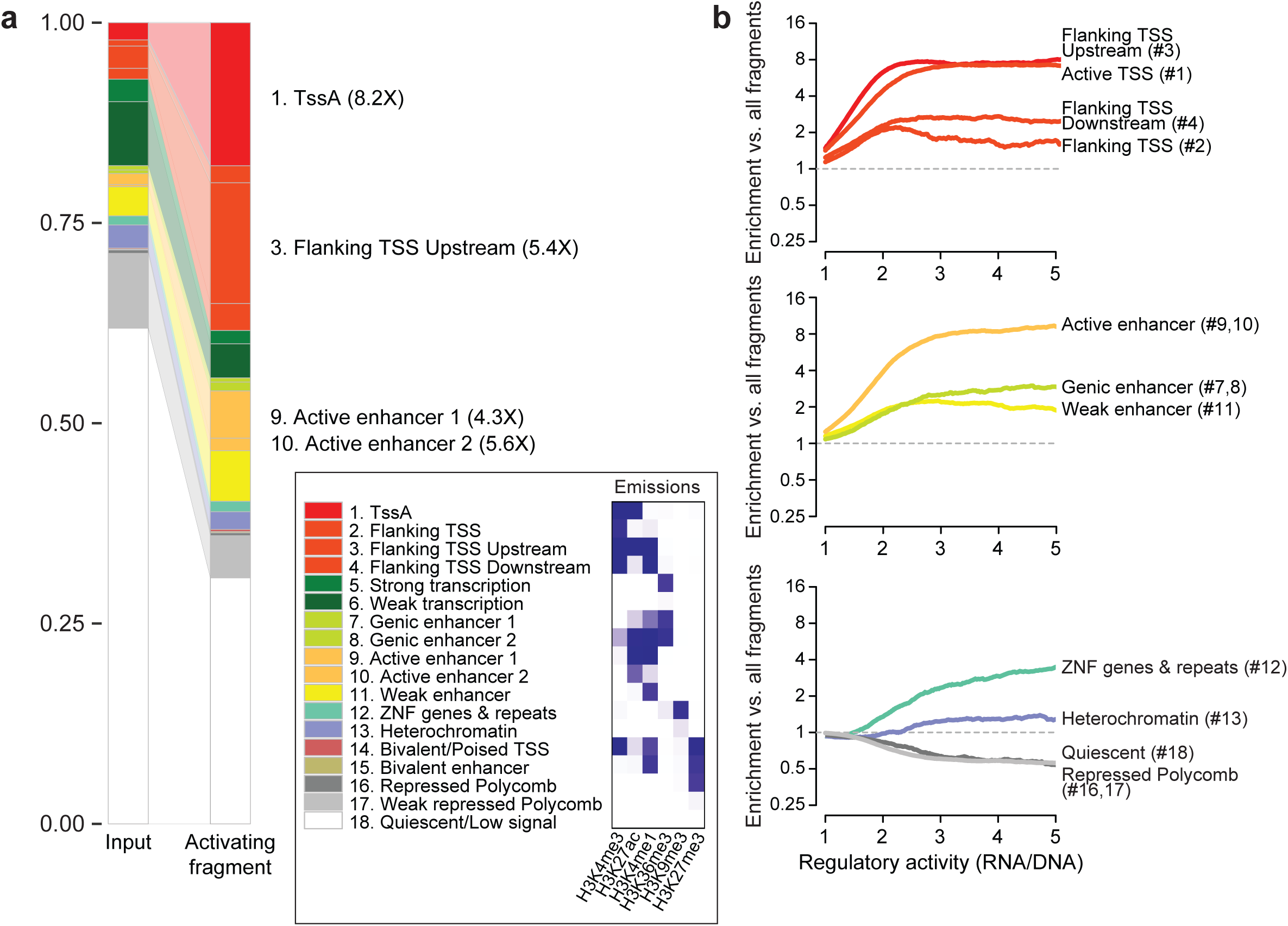
Active HiDRA fragments are enriched in endogenously active regulatory regions. (a) Overlap of active HiDRA fragments with different endogenous chromatin states. Heights correspond to proportion of nucleotides within active HiDRA fragments in each chromatin state. *Inset*: histone modification enrichments in each of 18 ChromHMM chromatin states (b) HiDRA fragment regulatory activity (fold-change increase in RNA levels) across different chromatin states. Numbers correspond to chromatin state numbers in 18-state ChromHMM model.

In addition to active promoter and active enhancer chromatin states, the “TSS Flanking Upstream” chromatin state showed strong enrichment for active HiDRA fragments (7.3-fold higher than expected from the input library). This chromatin state is defined by the presence of both promoter and enhancer histone marks H3K4me1, H3K4me3, and H3K27ac, and was named “TSS Flanking” due its depletion at exactly the TSS position, but its enrichment 400nt-1kb upstream of annotated transcription start sites^9^. However, 64% of its occurrences are >2kb from the nearest transcription start site, suggesting that a portion of genomic regions annotated as “TSS Flanking Upstream” may function biologically as distal enhancers (**Fig. 4a**).

**Figure 4:**
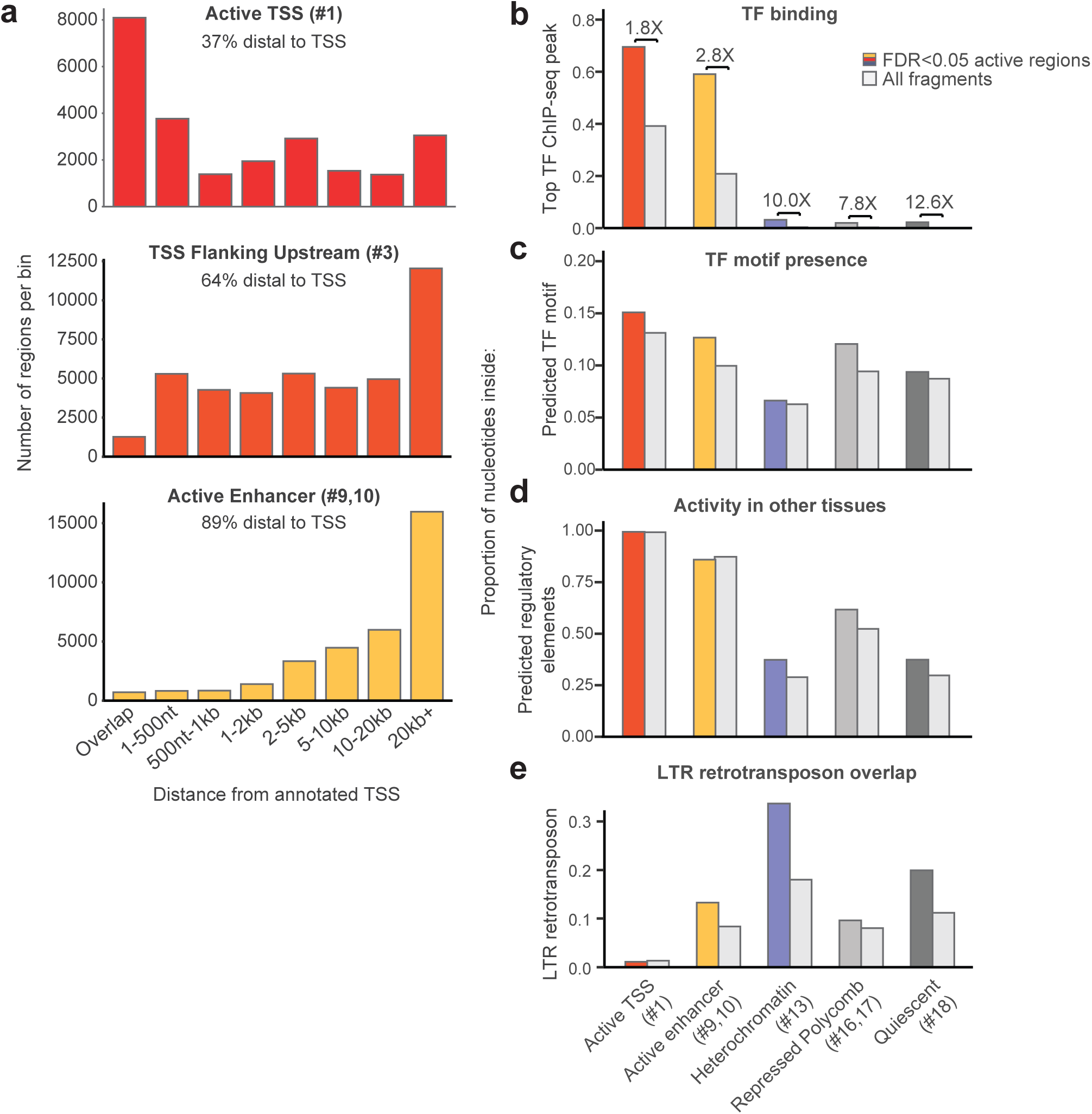
HiDRA activity outside of promoter and enhancer elements.

When we computed enrichment of chromatin states as a function of HiDRA activity strength, we found a linear quantitative relationship for HiDRA activity levels up to ~2.5-fold RNA/DNA ratios, with increasing activity showing increasing chromatin state enrichment for both promoter and enhancer chromatin states (**Fig. 3b**). Surprisingly, this enrichment stayed constant thereafter for promoter regions, and increased modestly for enhancer regions, ultimately surpassing the enrichment seen for promoters. In fact, even though promoter chromatin states were more enriched at intermediate HiDRA activity levels, enhancer chromatin states were the most enriched at the highest HiDRA activity levels (p=9.3x10^−102^, Supplemental **Fig. S3a**), suggesting that enhancer elements have a greater dynamic range of regulatory activity potential, which has implications for the regulatory architecture of genes.

At the other end of the spectrum, Quiescent and Polycomb-repressed chromatin states showed a 2-fold depletion for HiDRA active elements, but heterochromatin-associated chromatin states showed a modest enrichment, indicating that they may contain regulatory signals that become active once taken outside their repressive endogenous chromosomal context. The ZNF/repeats-associated chromatin state (marked by H3K36me3 and H3K9me3) showed a modest enrichment for lower HiDRA activity levels, but continued to increase linearly even at the highest activity levels, possibly due to active repetitive elements, as we discuss below.

We also studied the enrichment of HiDRA regions for individual histone marks profiled by the ENCODE project in GM12878^9^. Active promoter- and active-enhancer-associated acetylation marks H3K9ac and H3K27ac, histone turnover-associated H2A.Z, promoter- and enhancer-associated H3K4me3 and H3K4me1, and DNase I accessible chromatin were the most enriched individual marks within active HiDRA regions, while Polycomb-repression-associated H3K27me3, heterochromatin-associated H3K9me3, and transcription-associated H3K36me3 were the least enriched compared to the input library (Supplemental **Fig. S4**).

As these elements are tested outside their endogenous chromatin context, we expect that they drive reporter gene transcription by recruiting transcriptional regulators in a sequence-specific way, and we sought to gain insights into the recruited factors. We calculated the overrepresentation of 651 transcription factor sequence motifs assembled by ENCODE in active HiDRA regions, and found enrichment for many distinct motifs for immune transcription factors (Supplemental **Fig. S3b**), including IRF, NFKB1, and RELA, corresponding to transcriptional regulators known to function in GM12878 compared to other human cell lines. The motifs enriched in promoter chromatin states were largely distinct from those enriched in enhancer chromatin states, highlighting the differential regulatory control of the two types of regions (Supplemental **Fig. S3a**). These differences in motif content indicate that the two types of regions recruit different sets of transcriptional regulators both in their endogenous context and in our episomal assays, consistent with their distinct endogenous chromatin state and their distinct properties in our HiDRA assays.

### HiDRA regulatory activity outside promoter and enhancer regions

Even though HiDRA active regions were most enriched for enhancer and promoter states, they were not exclusive to them. In fact, approximately half of active HiDRA regions (52%) showed endogenous epigenomic signatures characteristic of repressed and inactive chromatin states, including Quiescent, Repressed Polycomb, Weak Repressed Polycomb, and Heterochromatin.

As active chromatin states were defined based on the profiling of only a subset of known chromatin marks in GM12878, we reasoned that perhaps other marks may be marking these regions active, but that they were perhaps not profiled in GM12878 and thus missed by the reference genome annotations. For example, a recent study identified subclasses of active enhancer elements marked with H3K122ac or H3K64ac but not H3K27ac^14^. While these marks were not profiled in GM12878, inactive chromatin states that showed HiDRA activity were 8-fold to 13-fold more likely to be bound by transcription factors in ChIP-seq experiments in GM12878 than inactive chromatin states that lacked HiDRA activity (**Fig. 4b**), indicating that our assays can successfully recover active regions even outside active chromatin states, and highlighting the importance of our unbiased survey of open chromatin regions regardless of their endogenous chromatin marks.

As both high-throughput and low-throughput episomal assays test regions outside their endogenous chromatin context, we reasoned that some active HiDRA regions with inactive chromatin signatures may reflect endogenously-inactive regions that become active when removed from the influence of nearby repressive effects. We reasoned that these regions would contain sequence motifs of TFs active in GM12878, but that these sequence motifs would be less likely to be bound *in vivo*, compared to motifs in active states. Indeed, we found that active HiDRA regions from endogenously-inactive chromatin states showed similar enrichments in regulatory motif occupancy to that of enhancer and promoter chromatin states (**Fig. 4c**), but substantial differences in their endogenous TF binding (**Fig. 4b**), consistent with endogenous repression due to their genomic context. These regions were also ~30% more likely to be active in another human tissue, compared to HiDRA-inactive regions (**Fig. 4d**), consistent with cell-type-specific repression in their endogenous chromatin context.

In addition to the presence of regulatory motifs for known regulators active in GM12878, we sought additional driver elements that may be responsible for the episomal activity of endogenously-inactive regions. In particular, we considered the presence of Long-Terminal-Repeat (LTR) retrotransposons, which have been previously shown to have regulatory activity potential and were enriched in the set of all active HiDRA regions unlike other repetitive elements in the genome (Supplemental **Fig. S5**)^2,20^. Indeed, we found that active HiDRA regions from endogenously-inactive regions showed substantial enrichment for LTR retrotransposons. In fact, Quiescent and Heterochromatin states were more enriched for LTR retrotransposons than either Enhancer or Promoter chromatin states (**Fig. 4e**, Supplemental **Fig. S5**). As LTRs are motif-rich and often act as the substrate for recently evolved enhancers, these endogenously-inactive but episomally-active HiDRA regions may represent a reservoir for the emergence of new regulatory elements^6^.

### High-resolution mapping of regulatory activity with HiDRA

We next sought to exploit the highly overlapping nature of tested HiDRA fragments to increase the resolution of regulatory inferences by exploiting subtle differences between neighboring fragments that only overlap partially. As an example, we considered a 3kb region on chromosome 7 that is covered by 134 distinct HiDRA fragments. When we examine every fragment in this region, we observed that fragments overlapping the known RUNX3 motif showed substantially higher regulatory activity (**Fig. 5a**). This motif is bound by the RUNX3 protein in GM12878 cells and shows increased evolutionary conservation (**Fig. 5a**). These properties suggest that the driver regulatory nucleotides within this region are tightly concentrated surrounding the RUNX3 motif, and that on the global level the differential activity of HiDRA-tested segments should enable us to systematically discover these driver nucleotides in an unbiased way based on the relative activity of fragments that do or do not overlap them.

**FIgure 5:**
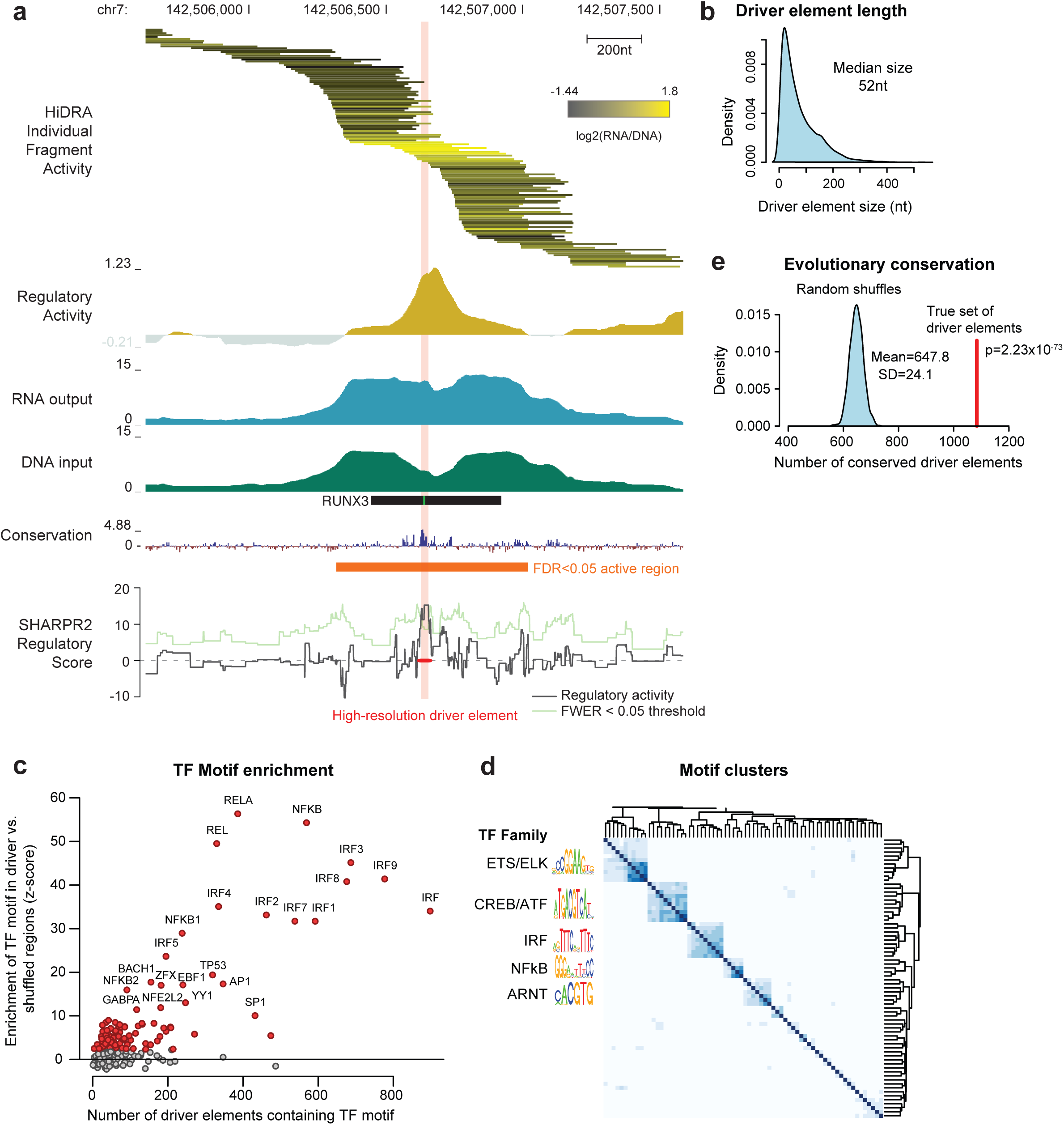
High-resolution mapping of transcriptional regulatory elements with SHARPR2. (a) Example region used in high-reso-tion mapping. Fragment activity shown on log2 scale with two fragments with highest and lowest activity removed for color scale to avoid outliers. The transparent red bar indicates the driver element identified at the regional FWER<0.05. (b) Size distribution of driver ements. (c) Enrichment of immune-related TF motifs in driver elements compared to shuffled driver elements within tiled regions. (d) TF motifs enriched in driver elements cluster into groups of co-occurring motifs, suggesting diversity of TF motifs involved in transcripnal regulatory activity (e) Significantly more driver elements are evolutionary conserved compared to shuffled driver elements within ed regions. Evolutionary conservation cut-off chosen as conservation score for top 5% of shuffled regions.

As part of our development of Sharpr-MPRA^2^, we had previously developed the SHARPR algorithm, a graphical probabilistic model that inferred high-resolution activity from MPRA tiling experiments by reasoning about the differential activity of partially-overlapping microarray spots. Intuitively, SHARPR allowed us to transform measurements from the 145-bp resolution of individually tested tiles to the 5-bp resolution of the offset between consecutive tiles. The SHARPR algorithm relies on synthesized oligos that uniformly tile regions at regularly spaced intervals, and thus is not applicable for the random fragmentation nature of HiDRA experiments where both the length and the spacing of neighboring fragments can vary. To address this challenge, we developed a new algorithm, SHARPR2, which estimates regulatory scores underlying any set of randomly-positioned and variable-length segments, by appropriately scaling the segments by their varying lengths, and enabling inferences at variable-length offsets between them (Supplemental Information).

Applying the SHARPR2 algorithm to the RUNX3 example above, we found that the 3kb region was narrowed down to a single ‘driver’ element of 27nt (**Fig. 5a**). These captured the known RUNX3 motif shown experimentally by ChIP-Seq to be bound by the RUNX3 regulator in GM12878^9^, and also the independently-determined high-resolution region of evolutionary conservation, even though neither line of evidence was used in our inferences.

Across all ~32,000 “tiled regions” that are covered by at least 10 unique HiDRA fragments, SHARPR2 predicted ~13,000 driver elements of median length 52nt, using a regional family-wise error rate of 5% (**Fig. 5b**). With increasing coverage, the resolution of driver regions also increased, from ~50nt for regions with 10-20 fragments to ~20nt for regions with 40 or more fragments (Supplemental **Fig. S6**). The length of driver elements did not further decrease between 40 and 80 fragments per tiled region, suggesting a minimum number of ~18 driver nucleotides necessary to drive regulatory activity. Similar to active HiDRA regions, driver elements were also mostly distal from annotated TSS regions, and were preferentially found in endogenously active chromatin states (active promoters, TSS-flanking, and active enhancer regions, Supplemental **Fig. S7**).

We found that predicted driver nucleotides were significantly more enriched for regulatory motifs than shuffled controls (obtained by randomly shuffling driver positions within tiled regions). The enriched motifs consisted of regulators known to be active in GM12878, including several critical B-cell and immune transcription factor including NF-kB and the IRF family (**Fig. 5c**). A total of 98 motifs were enriched in driver elements (FDR<0.05), clustering into several distinct groups with little overlap between groups, suggesting a wide range of distinct transcription factors act to regulate GM12878 gene expression (**Fig. 5d**). We also found that driver nucleotides are significantly more likely to be evolutionarily-conserved across vertebrates than randomly-shuffled controls (**Fig. 5e**), with ~1080 driver elements overlapping conserved regions, compared to only ~650 expected by chance (p=2.23x10^−73^). These results indicate that our high-resolution inferences are biologically meaningful and can help pinpoint driver nucleotides among larger regions.

### Prioritization and characterization of GWAS variants affecting regulatory activity

We next sought to use our predicted active regions and driver nucleotides to gain insights into non-coding variation. We studied the overlap between genetic variants associated with immune disorders and our high-resolution predicted driver nucleotides. Even though driver nucleotides only cover 0.032% of the genome, we found 12 cases where they overlap fine-mapped SNPs associated with 21 immune-related traits^21^ predicted to be causal (~5 expected by chance inside tiled regions, p=0.012, **Fig. 6a**). For example, we predict a 76-nt driver element overlapping rs12946510 in the IKZF3 locus associated with multiple sclerosis in a tiled region of 3kb (**Fig. 6b**), suggesting this may be the causal variant. The SNP overlaps a 76-nt driver element that contains a RUNX3 motif and a RELA motif, both bound by the respective TFs in GM12878^9^.Indeed, rs12946510 is predicted to be causal based on genetic fine-mapping^21^, with a posterior probability of 0.314 of being causal with the next strongest signal showing only 0.067 posterior probability. rs12946510 is also an eQTL for the IKZF3 gene^21,22^, and was recently shown to disrupt enhancer activity for the surrounding 279-nt region using a luciferase reporter assay^23^, consistent with our prediction that rs12946510 is a causal SNP.

**Figure 6:**
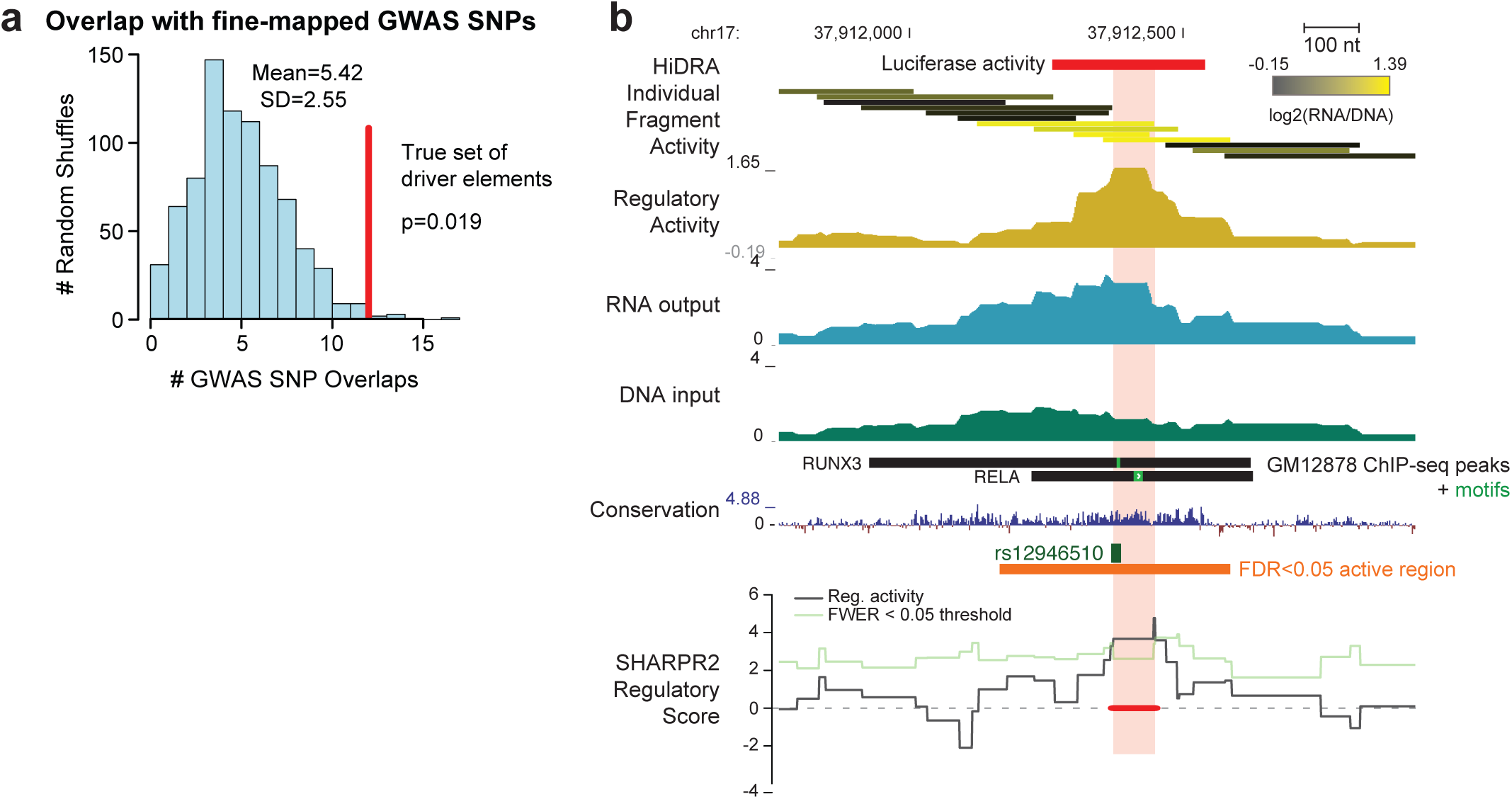
High-resolution driver elements are enriched for fine-mapped GWAS SNPs. (a) Driver elements overlap more GWAS fine-mapped SNPs associated with 21 human immune-related complex traits than randomly shuffled regions. (b)Example locus at rs12946510 that overlaps a high-resolution driver element. Highlighted segment indicates the driver element identified at the regional FWER<0.05. Red bar at top corresponds to region with luciferase activity as demonstrated by Hitomi etal. (2017).

To recognize regions that showed differential activity between risk and non-risk alleles of common genetic variants, we first inferred the genotype of all RNA fragments profiled. As HiDRA is a sequencing-based assay, where the expression of reporter genes is quantified based on the number of sequencing reads, allele-specific differences in HiDRA activity between risk and non-risk haplotypes should be detectable in principle by using heterozygous positions to distinguish reads coming from the paternal or the maternal allele. In practice however, HiDRA fragments are much longer (~337 median length) than the typical sequencing reads we used for quantification (37nt, paired end), and thus 78% of genetic variants will not be covered by our sequencing reads (if they fall in the inner ~260nt not captured by our paired-end sequencing). To overcome this limitation and to determine allele-specific activity scores for all our fragments, we used low-depth re-sequencing of our input library using long reads, thus revealing the genotype associated with each start/end position in our library (**Fig. 7a**). We augmented this information with 4-nt random i7 barcodes that were added by PCR during the initial HiDRA library construction, thus ensuring that the [*start*, *end*, *i7*] triplet is almost guaranteed to be unique, by resolving the cases where both start and end positions are identical between paternal and maternal alleles. This strategy enabled us to resolve the genotype of all previously quantified HiDRA fragments without having to sequence both the plasmid and RNA libraries to full length at high depth, which would be too costly.

**Figure 7:**
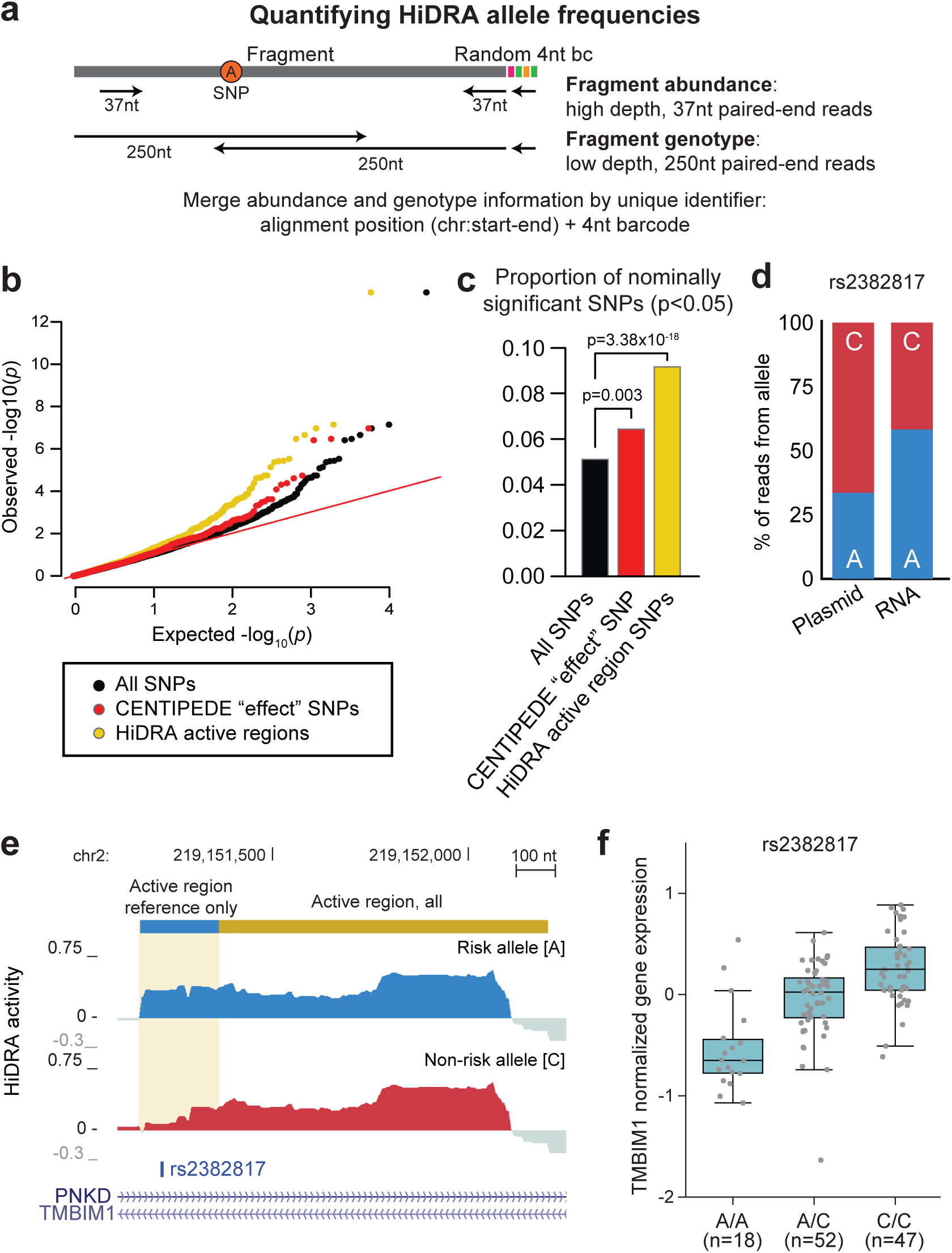
Identification of human genetic variants that alter HiDRA activity. (a) Overview of genotyping approach for HiDRA fragments. HiDRA fragments were originally quantified at high-depth using 37nt paired-end reads. At this read length the allele composition of fragments is mostly unobserved. As every HiDRA fragment has a unique identifier (genomic alignment position and random 4nt barcode), long-read re-sequencing of the HiDRA library can assign SNP genotypes to fragments that were previously quantified for activity using short reads. (b) q-q plot for allelic imbalance at SNPs covered by HiDRA fragments. CENTIPEDE “effect” SNPs were identified by Moyerbrailean *et al.* (2016). (c) “Effect” SNPs and SNPs within HiDRA active regions are more likely to be nominally significant for allelic imbalance. (d) The A allele of rs2382817, a SNP associated with inflammatory bowel disease, is more active in the HiDRA assay than the C allele. (e) Alelle-specific HiDRA activity signal tracks for rs2382817. (f) rs2382817 alleles are correlated with differences in expression of the nearby TMBIM1 gene in EBV-transformed lymphocytes.

In a proof-of-concept analysis to assess the ability of HiDRA to detect allelic activity, we applied this approach systematically to all heterozygous positions known in the genotyped GM12878 cell lines. We found ~180,000 heterozygous SNPs that were represented by at least one HiDRA fragment at either allele in our library. We realized that fragments carrying the maternal or paternal alleles of a SNP may also differ at their start and end positions, and that differences at fragment ends may cause SNPs with no true biological activity to falsely appear to disrupt HiDRA activity (Supplemental **Fig. S8**). We attempted to filter out these cases by only comparing fragments that show 90% mutual overlap, and where the start and end of the fragment is more than 25nt from a high-resolution driver element, thus ensuring that allelic differences are not due to differential inclusion of driver elements (~16,000 SNPs remained after filtering). At an uncorrected nominal p-value cut-off of 0.05, we found 880 ‘allelic’ HiDRA SNPs where paternal and maternal alleles showed differences in activity, 25 of which had a corrected FDR<0.1 (beta-binomial model^24^). The corresponding SNPs in these 880 allelic HiDRA regions were more frequently found in HiDRA active regions and more frequently predicted to have strong regulatory effects in open chromatin regions by an independent study^25^ (**Fig. 7b,c**), suggesting they are biologically meaningful.

For example, we found that rs2382817, a SNP associated with inflammatory bowel disease^22^ (p_GWAS_=1.13x10^−13^), shows differential HiDRA activity between paternal and maternal alleles. The risk allele shows increased regulatory activity upstream of a HiDRA-annotated active region (nominal p=8.7x10^−4^, FDR=0.25, **Fig. 7d,e**). In a panel of human individuals from the GTEx project^26^, rs2382817 was an eQTL for TMBIM1 in EBV-transformed lymphocytes (the same cell type as GM12878, **Fig. 7f**), and for TMBIM1 and other nearby genes PNKD, ARPC2 and GPBAR1 in other tissues, consistent with a role of rs2382817 in gene expression regulation, and illustrating the possibility of using HiDRA to detect SNPs with allelic effects on regulatory activity.

These results indicate that HiDRA can help shed light on disease-associated variants, by either narrowing down the set of candidate causal SNPs using our high-resolution driver nucleotide inferences, or by directly observing differential activity between risk and non-risk alleles using allele-specific activity inferences.

## Discussion

We presented a high-throughput experimental assay, HiDRA, to test transcriptional regulatory activity for millions of DNA fragments preferentially generated from regions of open chromatin and discover high-resolution driver elements. We performed HiDRA mapping of regulatory activity using a library of sequences from the GM12878 lymphoblastoid cell line ranging from 169-477nt in length. We found that the endogenous loci of up-regulated HiDRA fragments are significantly more likely to be classified as promoter and enhancer elements, contain motifs for immune transcription factors and be marked by activating histone modifications. We also leverage the dense tiling of HiDRA fragments at regulatory regions to perform ahigh-resolution mapping of regulatory activity to identify short DNA segments that act as drivers of regulatory activity, including one 76nt driver element that overlaps a SNP, rs12946510, associated with multiple sclerosis risk.

While we performed our study in the GM12878 cell line, the HiDRA methodology can be readily applied to study the transcriptional regulatory architecture of any cell line. For cell lines with poor transfection efficiencies, a non-integrating lentiviral infection method can be used instead of transfection, as both approaches have shown highly similar results in other high-throughput reporter assays^27^. HiDRA libraries can also be transfected in a different cell line than was used for library generation. For example, libraries could be generated from limited patient tissue, and subsequently transfected into a relevant immortalized cell line.

We also demonstrate a proof-of-concept application of HiDRA to identify SNPs that alter regulatory element activity by mapping reads in an allele-specific manner, as well as illustrate potential confounding factors for analyzing allelic HiDRA and STARR-seq data when fragments mapping to either allele have different genomic positions. Moving forward, construction of libraries with higher coverage at relevant SNPs should mitigate this concern. HiDRA also relies on the presence of different alleles in the input genomic DNA. While no human individual or cell line exists that is heterozygous at every clinically important genetic variant, future studies can pool cells or tissue from multiple individuals to generate a HiDRA library heterozygous at more loci. As recent studies have also shown that cancer driver mutations are enriched inside promoter elements, HiDRA may also be applied to pools of tumor samples to identify promoter variants that experimentally alter regulatory activity and gene expression^28^.

One limitation of HiDRA is the use of genomic DNA, while technologies involving *in vitro* synthesis can readily introduce changes to DNA not observed in the human population to better fine-map regulatory sub-regions of enhancers^15,29^. HiDRA fragment libraries could likewise be modified to introduce non-existing mutations through error-prone PCR or introduction of mutagens during fragment amplification. Another potential improvement to the assay might be to further enrich for fragments from active regulatory regions, by coupling with a fragment capture technology similar to those used in Capture Hi-C to selectively test a subset of enhancers or promoters at higher resolution while retaining the advantages of having larger fragment sizes and high library complexity^30^. Finally, the SHARPR2 high-resolution mapping algorithm can be applied to other STARR-seq experiments. For example, investigators interested in only a specific locus could perform STARR-seq on a bacterial artificial chromosome clone (“BAC-STARR-seq”) that contains the region of interest^3^. SHARPR2 high-resolution mapping will then readily be capable of mapping regulatory activity of this specific locus with single nucleotide resolution.

In summary, we present HiDRA, a high-throughput method to assay the regulatory activity of millions of open chromatin-derived fragments located genome-wide. As HiDRA can be readily applied to any eukaryotic cell type, we envision this approach or similar technologies being used to quantify the transcriptional regulatory landscape of DNA sequences for a variety of tissues from multiple organisms.

## URLs and Data Availability

All high-throughput sequencing data generated by this study has been deposited in NCBI GEO with accession GSE104001. Processed HiDRA plasmid input, RNA output, activity, as well as active fragments and driver elements can be visualized on the UCSC genome browser at: https://genome.ucsc.edu/cgi-bin/hgTracks?hgS_doOtherUser=submit&hgS_otherUserName=xinchenw&hgS_otherUserSessionName=HiDRA_GM12878_092617. The SHARPR2 R package is currently available from R-forge at: https://r-forge.r-project.org/R/?group_id=2288, and will also be available from CRAN pending package approval.

## Acknowledgements

We wish to thank E. Oltz and Y. Huang (Washington U. in St. Louis) for sharing GM12878 luciferase data; Y. Park for helpful discussions for SHARPR2 model; R. Tewhey for helpful discussions and advice on library construction and experimental protocols and; E. Brown for advice on GM12878 transfections; N. Rinaldi, V. Subramanian and L. Surface for helpful comments on the manuscript. M.C. and M.K. were supported by NIH R01 HG008155.

## Online Methods

We strongly recommend reading the Supplemental Notes for additional information and considerations about the Methods.

### HiDRA library construction

We performed 16 ATAC-seq reactions on 50,000 GM12878 cells each using a modified protocol based upon Buenrostro *et al*. (Supplemental Note 1). We performed cell collection, lysis, and Tn5 digestion as described by Buenrostro *et al.*, Tn5-fragmented DNA was cleaned up using a MinElute PCR purification kit (Qiagen #28004, four reactions per column eluted in 20uL EB buffer) and the resulting 80uL of eluate was split into 16 PCR reactions (Supplemental Note 2). PCR was performed using custom HPLC-purified primers (F: 5’-TAGAGCATGCACCGGCAAGCAGAAGACGGCATACGAGATNNNNATGTCTCGTGGGCTCGGAGATGT-3’, R: 5’-GGCCGAATTCGTCGATCGTCGGCAGCGTCAGATGTG-3’, NNNN corresponds to random 4nt i7 barcode sequence) and NEBNext Ultra II Q5 DNA polymerase master mix (NEB #M0544L). Thermocycler conditions were: 65C for 5 min, 98C for 30 sec, 8 cycles of: 98C for 10 sec and 65C for 90sec. PCR reactions were pooled and cleaned up with a Qiagen MinElute PCR purification kit (two PCR reactions per column eluted in 20uL EB buffer) and run on a 1% agarose E-Gel EX with SYBR Gold II stain (Thermo Fisher #G402001). Size selection of ATAC-seq fragments was performed by gel excision using a razor blade. Gel slabs were pooled into <300mg groups and DNA was purified using a MinElute Gel Extraction kit (Qiagen #28604), and eluted in 20 uL of buffer EB per column following modified guidelines described in Box 2 of Taiwo *et al.* (2012) ^31^. The resulting size-selected ATAC-seq fragment library was treated with an anti-mitochondrial DNA CRISPR/Cas9 library following the protocol outlined in Montefiori *et al.* using 10X excess of Cas9 protein (Supplemental Note 3) ^32^. We cleaned up the reaction with a Qiagen MinElute PCR purification kit and split into 8 PCR reactions for a second round of PCR using the same conditions and primers described above.PCR products were cleaned up using two rounds of AMPure bead selection (0.8X ratio of beads to input) to size-select against small fragments, eluted in 40uL of dH2O and quantified using a Qubit dsDNA HS Assay kit (Thermo Fisher #Q32854).

The pSTARR-seq_human plasmid used for generating the plasmid library was a gift from Alexander Stark (Addgene plasmid #71509). The linear backbone used for the subsequent cloning steps was generated by digesting 4ug of circular pSTARR-seq_human for 4-6 hours with AgeI and SalI restriction enzymes (NEB #R3552S and R3138S), followed by gel excision under a dark reader transilluminator (Clare Chemical #DR22A) to extract a linear 3.5kb fragment corresponding to the human STARR-seq plasmid backbone. We performed cloning of the fragment library into the plasmid backbone approximately following the Methods section from Arnold *et al.* (2013)^3^. For each library, we performed 20 individual InFusion HD cloning reactions (Takara Bio #638911) using a 3.5:1 molar ratio of insert to vector backbone, following manufacturer’s instructions (Supplemental Note 4). Each group of five InFusion reactions was collected and cleaned up using the Qiagen MinElute Enzymatic Reaction cleanup kit, eluted in 10uL of dH2O, and transformed into four 20uL aliquots of MegaX DH10B T1R electrocompetent bacteria. The bacteria were thawed on ice for 10 min and mixed with eluted DNA (five InFusion reactions per 100uL of bacteria). 22uL of bacteria/DNA mixture were pipetted into a 0.1cm electroporation cuvette (Thermo Fisher Scientific #P41050) and tapped repeatedly against a hard surface to remove bubbles. Cuvettes were electroporated using a Bio-Rad Gene Pulser Xcell Microbial Electroporation System (Bio-Rad #1652662) using the conditions: 2.0 kV, 200 ω, 25 μF (Supplemental Note 5). For high-yield transformations, we observed electroporation time constants between 4.8 and 5.1ms. After electroporation, bacteria were immediately collected in 750uL pre-warmed SOC media, pooled, and incubated for 1hr in a 37C shaker. After recovery, serial dilutions of bacteria were plated to estimate the number of clones in the library. Recovered bacteria were diluted in 2L of pre-warmed luria broth and 100ug/mL of carbenicillin and grown overnight (8-10 hours while shaking). Plasmids were collected from bacteria using the Plasmid Plus MegaPrep kit (Qiagen#12981) following manufacturer’s instructions. Plasmid concentration was quantified using a Nanodrop One machine (Thermo Scientific) and diluted to a 3ug/uL concentration for subsequent transfection steps. To ensure plasmid library quality and diversity, a small aliquot of the fragment library was amplified by PCR using i5 and i7 primers, run on an Illumina MiSeq machine using the 50-cycle v2 kit as per manufacturer’s instructions, and aligned to the human genome to ensure correct complexity and sufficient proportions of reads within predicted transcriptional regulatory elements (Supplemental Note 6, see subsequent Methods sections for details on processing of sequencing libraries).

### Cell culture and transfections

GM12878 cells were obtained from the Coriell biorepository and grown in RPMI 1640 Medium with GlutaMAX Supplement (Thermo Fisher #61870127), 15% fetal bovine serum (Sigma Aldrich #F2442), and 1% pen/strep at a density of between 2x10^5^ and 1x10^6^ cells/mL with regular media changes every 2-3 days. Approximately 24 hours before transfection, GM12878 cells were split to a density of 4x10^5^ cells/mL to ensure the presence of actively dividing cells for increased transfection efficiency. For transfection, cells were collected by centrifugation for 5 min at 300g, washed once with pre-warmed PBS, and collected again for 5 min at 300g. PBS was aspirated and cell pellets were re-suspended in Resuspension Buffer R (Thermo Fisher Scientific #MPK10096) at a concentration of 7.5 million cells per 100uL. DNA was added to cells at a concentration of 5ug of plasmid per 1 million cells. In total, we transfected between 120-130M million cells per replicate using 100uL tips from the Neon Transfection System at 1200V with 3 pulses of 20ms. Transfected cells were immediately recovered in pre-warmed GM12878 media without antibiotic and recovered at a density of 1x10^6^ cells/mL for 24 hours. In parallel, we performed two transfections of GM12878 cells with a positive control GFP plasmid to assess transfection efficiency using the same conditions.

### RNA isolation and cDNA generation

GM12878 cells were collected 24 hours post-transfection, washed twice in pre-chilled PBS (collecting for 5 min at 300g) and RNA was purified using the Qiagen RNEasy Maxi kit (Qiagen #75162) following manufacturer’s instructions and performing the optional on-column DNase treatment step (Qiagen #79254). Poly A+ RNA was extracted from total RNA using the Oligotex mRNA Midi kit (Qiagen #70042, two columns per RNA sample), and any remaining DNA was digested with a second DNase treatment step using Turbo DNase (Thermo Fisher #AM2238) following manufacturer’s instructions (Supplemental Note 7). Treated mRNA was cleaned up and concentrated using the Qiagen RNEasy MinElute Cleanup kit (Qiagen #74204). We generated cDNA from mRNA using Superscript III Reverse Transcriptase (Thermo Fisher #18080085) with a gene-specific RT primer located in the 3’UTR of the sgGFP reporter gene downstream from the inserted fragments (5’-CAAACTCATCAATGTATCTTATCATG-3’). Reverse transcription was performed following manufacturer’s recommendations except with 2ug of poly A+ mRNA and 1uL of 12.5uM primer per 20uL reaction, and extension was performed for 60 minutes at 50C (Supplemental Note 8). Reverse transcription reactions were cleaned up using a MinElute PCR purification kit (Qiagen #28106, two reactions per column) and eluted in 15uL of pre-warmed buffer EB.

### Library construction and high-throughput sequencing

We performed a qPCR to test the number of cycles needed for amplification of single-stranded cDNA as well as input material of plasmid DNA needed such that both reactions had the same Ct values. We used 1uL of ssDNA and dilutions of plasmid DNA similar to the method described by Tewhey *et al.* Cell 2016. qPCRs were performed in 10uL reactions with all reagents scaled down proportionally from a normal 50uL PCR reaction (1uL of DNA, 5uL of Ultra II Q5 master mix, 0.4uL of 25uM primer mix, 0.2uL of 10X SYBR dye, 3.4uL of dH2O) with thermocycler conditions: 98C for 30s, 20 cycles of: 98C for 10s, 65C for 90s. We proceeded to perform eight regular 50uL PCR reactions (each scaled up 5X from the 10uL PCR reactions) using the same thermocycler conditions except using the Ct value for the cycle number (F: 5’-CAAGCAGAAGACGGCATACGAGAT-3’, R: 5’-AATGATACGGCGACCACCGAGATCTACAC[X8]TCGTCGGCAGCGTC-3’, “X8” sequence corresponds to sample barcode, chosen from Illumina Nextera barcode list). PCR reactions were cleaned up using Qiagen MinElute PCR purification kits and balanced for sequencing using the Kapa Library Quantification Kit (Kapa Biosystems #KK4824, Supplemental Note 9).

Each library batch (five transfected RNA biological replicates, five plasmid controls) was sequenced by the Broad Institute Walk-Up Sequencing Facility on four flowcells on a NextSeq 500 machine using the 75-cycle kit as per manufacturer’s instructions for 2x37nt paired-end reads with 2x8nt barcodes.

### Read mapping, data processing and identification of significantly up-regulated fragment groups

Reads were labelled by a random 4nt P7 barcode and an 8nt P5 barcode for sample ID. Reads were split into the ten samples (5 plasmid replicates and 5 RNA replicates) by P5 barcode and aligned to the human genome (hg19 assembly) using bowtie2 v2.2.9. Alignment files were filtered to (i) keep only aligned fragments, (ii) remove reads mapping to chrM, (iii) select reads passing the -q 30 filter in samtools, and (iv) remove reads aligning to the ENCODE hg19 blacklist regions (Supplemental Note 10). We identified unique fragments using the bamtobed command in BEDTools (v2.26.0) and filtered to keep only fragments between 100 and 600nt.

In analyzing results from HiDRA, we track the abundance of each individual fragment between the input (plasmid DNA) and output (RNA). We grouped fragments into “fragment groups” by 75% mutual overlap (bedtools v2.26.0), removed redundant fragment groups and summed counts of all fragments per group. To control for length-dependent biases, we split fragment groups into bins of 100nt (100-200, 200-300, etc.) and used DEseq2 (v1.10.1) to identify FDR<0.05 significantly up-regulated fragment groups^33^.

### Analysis of active HiDRA regions

All overlap and shuffle analyses performed using the BEDTools suite, v2.26.0^34^. Most colors for plots chosen with guidance from the wesanderson R package (https://github.com/karthik/wesanderson). For chromatin state annotations we used the 18-state output model generated by the Roadmap Epigenomics Consortium^1^. Active enhancer states were merged from states #9 and #10 (EnhA1 and EnhA2). ATAC-seq peaks positions were obtained from Buenrostro *et al.* (2013)^4^.

#### Signal tracks

Signal tracks for regulatory activity calculated as (RNA-DNA)/DNA after adding a pseudocount of 0.1 to both plasmid and RNA samples, and drawn in UCSC Genome browser showing only means (no whiskers) and with 5-pixel smoothing.

#### Correlation between RNA samples

We show correlations for fragments selected by four different cut-offs of minimum RPM. Pearson and Spearman correlations were calculated on log2-transformed data. Matrix of graphs drawn using layout and grid.arrange functions in R from the *gridExtra* library. Scatterplot between RNA samples drawn using the hexbinplot function from the *hexbin* library in R with xbins=100.

#### Proximal vs. distal

TSS regions were defined using the UCSC Genome Browser’s Table Browser tool for hg19. Distances to nearest annotated TSS were taken using closestBed tool in the BEDTools2 suite.

#### TF motif enrichment

We obtained the hg19 TF motif catalog from the ENCODE project^9^. We only considered motifs corresponding to transcription factors expressed in GM12878 (RPKM>5 using processed RNA-seq data from the Roadmap Epigenomics Consortium).

#### Activity of HiDRA regions in other tissues

We set a lenient definition for active in other tissues as the union of regions annotated in 97 non-GM12878 tissues from epigenome roadmap predicted with 18-state ChromHMM model. For active regions we considered states “TssA” (state #1), “TssFlnkU” (state #3), and “EnhA” (states #9 and 10).

#### SHARPR2 activity plots

Tracks were drawn in the UCSC Genome Browser using “Custom Tracks”. Coloring of individual fragments was performed by setting maximum and minimum colors (RGB 0,0,0 and RGB 255,255,0, respectively) to log2(RNA/DNA) values of 3rd lowest and 3rd highest fragments (two strongest and weakest fragments were removed to avoid strong outliers), and scaling colors of all other fragments linearly between these extremes. We chose to include only ChIP-seq bound TF bars for ChIP-seq experiments performed in GM12878 cells by the ENCODE project and where the motif (green bar) overlapped driver nucleotides.

### SHARPR2 identification of high-resolution driver elements

See Supplemental Information for details and more information on SHARPR2.

### Read mapping and data analysis for allele-specific regulatory activity

We used vcf-consensus (VCFTools) to mask the hg19 genome assembly by replacing heterozygous nucleotides identified by the Illumina NA12878 Platinum Genome with N’s. 250nt paired-end MiSeq reads were trimmed using cutadapt to remove Illumina primer sequences, mapped to the NA12878-masked hg19 assembly using bowtie2 v2.2.9 (settings: --end-to-end --phred33 --sensitive -p 7 -N 1 --no-unal), and filtered using the steps described above for 37nt reads. As some long reads have poor quality scores at their 3’ end, we trimmed low quality sequences (quality value < 38) to reduce the proportion of sequencing errors at SNPs that could lead to incorrect allelic assignment of fragments. Fragments were then assigned to a SNP based on genotype at the position. For comparisons of SNP activity, we only considered fragments with 90% mutual overlap to reduce the confounding effect of fragments that differ by both allele and position. We also removed fragments if either end was within 25nt of a driver element, as in these cases small differences in end position could artifically lead to large effects. After assigning fragment abundances (from high-depth 37nt PE read sequencing) to each allele of a SNP, we identified SNPs with significant differential activity using QuASAR-MPRA. CENTIPEDE SNPs were identified by Moyerbrailean *et al.* (2016) using an effect-size cut-off of >3 or <-3, following the cut-offs used by Kalita *et al.* (2017)^24,25^.

## References

1. Consortium, R. E. et al. Integrative analysis of 111 reference human epigenomes. Nature 518, 317– 330 (2015).

2. Ernst, J. et al. Genome-scale high-resolution mapping of activating and repressive nucleotides in regulatory regions. Nat Biotechnol 34, 1180–1190 (2016).

3. Arnold, C. D. et al. Genome-wide quantitative enhancer activity maps identified by STARR-seq. Science 339, 1074–1077 (2013).

4. Buenrostro, J. D., Giresi, P. G., Zaba, L. C., Chang, H. Y. & Greenleaf, W. J. Transposition of native chromatin for fast and sensitive epigenomic profiling of open chromatin, DNA-binding proteins and nucleosome position. Nature Methods 10, 1213–1218 (2013).

5. Nord, A. S. et al. Rapid and Pervasive Changes in Genome-wide Enhancer Usage during Mammalian Development. Cell 155, 1521–1531 (2013).

6. Long, H. K., Prescott, S. L. & Wysocka, J. Ever-Changing Landscapes: Transcriptional Enhancers in Development and Evolution. Cell 167, 1170–1187 (2016).

7. Wang, X. et al. Discovery and validation of sub-threshold genome-wide association study loci using epigenomic signatures. eLife Sciences 5, e10557 (2016).

8. Ernst, J. & Kellis, M. Discovery and characterization of chromatin states for systematic annotation of the human genome. Nat Biotechnol 28, 817–825 (2010).

9. ENCODE Project Consortium et al. An integrated encyclopedia of DNA elements in the human genome. Nature 489, 57–74 (2012).

10. Rada-Iglesias, A. et al. A unique chromatin signature uncovers early developmental enhancers in humans. Nature 470, 279–283 (2010).

11. Ernst, J. et al. Mapping and analysis of chromatin state dynamics in nine human cell types. Nature473, 43–49 (2011).

12. Visel, A., Minovitsky, S., Dubchak, I. & Pennacchio, L. A. VISTA Enhancer Browser--a database of tissue-specific human enhancers. Nucleic Acids Res. 35, D88–D92 (2007).

13. Taylor, G. C. A., Eskeland, R., Hekimoglu-Balkan, B., Pradeepa, M. M. & Bickmore, W. A. H4K16 acetylation marks active genes and enhancers of embryonic stem cells, but does not alter chromatin compaction. Genome Res. 23, 2053–2065 (2013).

14. Pradeepa, M. M. et al. Histone H3 globular domain acetylation identifies a new class of enhancers. Nature Genetics 48, 681–686 (2016).

15. Tewhey, R. et al. Direct Identification of Hundreds of Expression-Modulating Variants using a Multiplexed Reporter Assay. Cell 165, 1519–1529 (2016).

16. Melnikov, A. et al. Systematic dissection and optimization of inducible enhancers in human cells using a massively parallel reporter assay. Nat Biotechnol 30, 271–277 (2012).

17. Kwasnieski, J. C., Mogno, I., Myers, C. A., Corbo, J. C. & Cohen, B. A. Complex effects of nucleotide variants in a mammalian cis-regulatory element. Proceedings of the National Academy of Sciences 109, 19498–19503 (2012).

18. Gillies, S. D., Morrison, S. L., Oi, V. T. & Tonegawa, S. A tissue-specific transcription enhancer element is located in the major intron of a rearranged immunoglobulin heavy chain gene. Cell 33, 717–728 (1983).

19. Huang, Y. et al. cis-Regulatory Circuits Regulating NEK6 Kinase Overexpression in Transformed B Cells Are Super-Enhancer Independent. Cell Reports 18, 2918–2931 (2017).

20. Barakat, T. S. et al. Functional dissection of the enhancer repertoire in human embryonic stem cells. biorxiv.org Available at: www.biorxiv.org/content/biorxiv/early/2017/07/04/146696.full.pdf.

21. Farh, K. K.-H. et al. Genetic and epigenetic fine mapping of causal autoimmune disease variants. Nature (2014). doi:10.1038/nature13835

22. Jostins, L. et al. Host–microbe interactions have shaped the genetic architecture of inflammatory bowel disease. Nature 491, 119–124 (2012).

23. Hitomi, Y. et al. Identification of the functional variant driving ORMDL3 and GSDMB expression in human chromosome 17q12-21 in primary biliary cholangitis. Sci Rep 7, 2904 (2017).

24. Kalita, C. A. et al. QuASAR-MPRA: Accurate allele-specific analysis for massively parallel reporter assays. biorxiv.org Available at: Error! Hyperlink reference not valid.

25. Moyerbrailean, G. A. et al. Which Genetics Variants in DNase-Seq Footprints Are More Likely to Alter Binding? PLoS Genet 12, e1005875 (2016).

26. GTEx Consortium. Human genomics. The Genotype-Tissue Expression (GTEx) pilot analysis: multitissue gene regulation in humans. Science 348, 648–660 (2015).

27. Inoue, F. et al. A systematic comparison reveals substantial differences in chromosomal versus episomal encoding of enhancer activity. Genome Res. 27, 38–52 (2017).

28. Khurana, E. et al. Role of non-coding sequence variants in cancer. Nat Rev Genet 17, 93–108 (2016).

29. Kheradpour, P. & Kellis, M. Systematic discovery and characterization of regulatory motifs in ENCODE TF binding experiments. Nucleic Acids Res. (2013). doi:10.1093/nar/gkt1249

30. Mifsud, B. et al. Mapping long-range promoter contacts in human cells with high-resolution capture Hi-C. Nature Genetics 47, 598–606 (2015).

31. Taiwo, O. et al. Methylome analysis using MeDIP-seq with low DNA concentrations. Nat Protoc 7, 617–636 (2012).

32. Montefiori, L. et al. Reducing mitochondrial reads in ATAC-seq using CRISPR/Cas9. Sci Rep 7, 1213 (2017).

33. Michael I Love, W. H. S. A. Moderated estimation of fold change and dispersion for RNA-seq data with DESeq2. Genome Biol 15, (2014).

34. Quinlan, A. R. & Hall, I. M. BEDTools: a flexible suite of utilities for comparing genomic features. Bioinformatics 26, 841–842 (2010).

